# Factors Associated with Suicidal Ideation and Suicidal Attempts among Adolescent Students in Nepal: Findings from Global School Based Students Health Survey

**DOI:** 10.1101/511105

**Authors:** Achyut Raj Pandey, Bihungum Bista, Raja Ram Dhungana, Krishna Kumar Aryal, Binaya Chalise, Meghnath Dhimal

## Abstract

**Introduction:** Suicide has been recognized as a public health problem with high burden in low and middle income countries. Suicide has long lasting psychological trauma on friends and relatives in addition to loss of economic productivity. Although the need of high quality evidence is acknowledged for designing suicide prevention program, Nepal lacks evidence from reliable and nationally representative data. In this context, this study aimed to estimate the prevalence of suicidal ideation and attempt and identify the factors associated with them.

**Materials and Methods:** Total of 6,531 students of grade 7 to 11 from 74 schools representing all three ecological belts and five development regions participated in this cross sectional study. To select the representative sample from study population, we used the two stage cluster sampling method. The respondents filled the self-administed standardized questionnaire. We carried out a Complex survey analysis for adjusting the selection probability of each research participants.

**Results:** Nearly 14% had considered suicide while 10.33% had attempted suicide. Factors associated with suicidal ideation were the food insecurity (OR=2.31, CI=1.64-3.27), anxiety (OR=2.53, CI=1.48-4.30), loneliness (OR=2.50, CI=1.41-4.44) and gender (OR=1.38, CI=1.03, 1.88) as risk factors of suicidal ideation and anxiety (OR=2.99, CI=1.18-7.64), loneliness (OR=2.19, CI=1.28-3.74) truancy(OR= 1.99, CI=1.39-2.87), cigarette use(OR=3.06, CI=1.32-7.09) and gender (OR=1.60, CI=1.07-2.39) as risk factors of suicidal attempt. Having 3 or more close friends was found to have protective effect (OR=0.35, CI=0.16-0.74) against suicidal attempt.

**Conclusions:** Study reveals the relatively high prevalence of suicidal ideation and suicidal attempt among school-going adolescents in Nepal. Appropriate coping strategies for factors like anxiety, loneliness seem could be useful for preventing both suicidal ideation and attempt.

## Introduction

Suicide is one of the major causes of death and disability worldwide. Annually, more than 800,000 people die of suicide, where a person commits suicide in every 40 seconds. [1]. The estimated age standardized rate for suicide was 10.67 per 100000 population in 2015 [2] disproportionately affecting young people of 15 and 29 years. Suicide is the second leading cause of death among 15–29-year-olds [3]. It is also largely distributed among low and middle-income countries, accounting almost 78% of all suicide deaths globally [3]. Suicide rate in Nepal is 7.2 in both sexed with 8.2 in male, and 6.3 in female per every 100000 population [1] [3]. Suicide in adolescence is often underreported with possible cause of death being classified as underdetermined or accident to protect the families from possible stigma associated with suicide [4].

Because of the gravity of issue, suicide has received attention in global public health arena in recent years. In addition to loss of life and economic productivity for the society, there is long lasting psychological trauma on friends and relatives. Prevention of suicide is the best option as most of suicide cases don’t get treatment or cannot be treated [5]. Target 3.2 of Mental Health Action Plan 2013-2020 has envisioned reducing the rate of suicide in WHO member countries by 10% by 2020 which requires clear understanding of the factors leading to suicide [6].

Although the information on epidemiology is essential for policy makers in designing interventions, suicidal behavior has a large number of underlying causes that are complex to understand and differ from one country to other thus making the preventive efforts more complex and diverse [7,8]. Both suicidal ideation and suicide share the common risk factors such as hopelessness, social isolation, anxiety, depression, impulsivity and substance abuse, among others. Several other adverse stimuli are also likely to be associated with the development of suicidal ideation and the progression from ideation to attempts [9]. Evidence suggests adolescents of both genders who had suicide thoughts and attempts are significantly more likely to commit suicide than those without such thoughts and attempts [10].

Delivery of mental health services including treatment of suicidal attempts are limited specially in city areas and is further constrained by limited human and financial resources with government spending being less than 1% of total healthcare budget in mental health. Thus suicide prevention program are crucial in context of Nepal. Suicide prevention activities need to be tailored as per the context of the country and need deeper understanding on the determinants of the suicide. However, Nepal lacks reliable and representative data due to multiple reasons like poor registration system, mis-categorization of suicide cases by hospitals, and underreporting of suicide incidences in police data because of stigma attached with suicide [11]. Although there are some data from hospital based studies confined to specific setting and small scale cross sectional studies, Nepal lacks large scale nationwide studies to guide policy making process [12,13]. Identification of factors associated with suicidal ideation among students can be helpful in designing the right interventions and reducing the burden of condition [14]. In this context, this study is designed to assess the determinants of suicide among adolescents of Nepal.

## Materials and Methods

Global School Based Students Health Survey (GSHS) 2015 was carried out using globally standardized methodology. Two-stage cluster sample was used in this study to select representative sample from study population that comprised of 7 to 11 class students. In first stage of sampling strategy, 74 schools were selected based on probability proportional to school enrolment size from among the total schools containing any of 7 to 11 class. In second stage, we randomly selected a intact classrooms from each of selected school to participate in the study and each student in the selected classroom were eligible to participate in the study. Varying probabilities of selection of each participants and non-response rate were adjusted applying appropriate weighting factor. Out of 74 schools selected for the study, 68 schools (92%) participated in the study. From among 8670 students selected for study, 6531 participated in the study making response rate of 75% while 6529 of the questionnaire were useable for the study making overall response rate of 69%.

Data were collected through self-administration of standardized questionnaire. The GSHS questionnaire used for this study contained 58 core questions, 33 expanded questions addressing all the GSHS core modules covering demographics of the students, dietary behaviors, hygiene, violence and unintentional injury, tobacco use, mental health, alcohol and drug use, sexual behaviors and physical activity. Variables used in this study, the survey questions used and their coding schemes are presented in table 1.

**Table 1:** Variables used in the study

| **Variable** | **Survey question** | **Coding** |
| --- | --- | --- |
| **Gender** | What is your sex? | 1=Male, 2=female |
| **Age** | How old are you? | 1=12 years or younger,<br>2=13 years, 3=14 years,<br>4=15 years, 5=16 years<br>or older |
| **Food insecurity** | During the past 30 days, how often did you go hungry because there was not enough food in your home? | 1=Most of times/always,<br>0=Never/rarely/sometime<br>s |
| **Anxiety** | During the past 12 months, how often have you been so worried about something that you could not sleep at night? | 1=Most of times/always,<br>0=Never/rarely/sometime<br>s |
| **Loneliness** | During the past 12 months, how often have you felt lonely? | 1=Most of times/always,<br>0=Never/rarely/sometime<br>s |
| **Close Friends** | How many close friends do you have? | 0=0 close friends , 1=1<br>close friends, 2=2 close<br>friends, 3=3 or more<br>close friends |
| **Truancy** | During the past 30 days, on how many days did you miss classes or school without permission? | 0=0 to 2 times, 1=3 or<br>more times |
| **Bullied** | During the past 30 days, on how many days were you bullied? | 0=0 times, 1=1 or more<br>times |
| **physically attacked** | During the past 12 months, how many times were you physically attacked? | 0=0 times, 1=1 or more times |
| **Physical fighting** | During the past 12 months, how many times were you in a physical fight? | 0=0 times, 1=1 or more times |
| **Current cigarette use** | During the past 30 days, on how many days did you smoke cigarettes? | 0=0 times, 1=1 or more times |
| **Initiation of drug use** | How old were you when you first used drugs? | 0= I have never used drugs, 1= any other response |
| **Really Drunk** | During your life, how many times did you drink so much alcohol that you were really drunk? | 0=0 times, 1=1 or more times |
| **Parents check homework** | During the past 30 days, how often did your parents or guardians check to see if your homework was done? | 1=Most of times/always,<br>0=Never/rarely/sometime<br>s |
| **Parents understand Problem** | During the past 30 days, how often did your parents or guardians understand your problems and worries? | 1=Most of times/always,<br>0=Never/rarely/sometime<br>s |
| **Parental monitoring** | During the past 30 days, how often did your parents or guardians really know what you were doing with your free time? | 1=Most of times/always,<br>0=Never/rarely/sometime<br>s |
| **Parental intrusion of privacy** | During the past 30 days, how often did your parents or guardians go through your things without your approval? | 1=Most of times/always,<br>0=Never/rarely/sometime<br>s |
| **Suicidal ideation** | During the past 12 months, did you ever seriously consider attempting suicide? | 0=No, 1=Yes |

| Variable | Survey question | Coding |
| --- | --- | --- |
| **Attempted suicide** | During the past 12 months, how many times did you actually attempt suicide? | 0=No, 1=Yes |

Data analysis was performed using STATA software version 15.0 (Stata Corporation, College Station, TX, USA). For all analysis, complex survey analysis was in the descriptive analysis, weighted percentages are reported. For inferential statistics, Chi square test and odds ratio (OR) was used for bivariate analysis to assess association between suicidal ideation/attempt and independent variables). All variables statistically significant at the *P* < 0.05 level in bivariate analyses were included in the multivariable models. Furthermore, multivarible analysis was used for evaluation of the effect of explanatory variables for suicide ideation/attempt in the past 12 months (binary dependent variables). The two-sided 95% confidence intervals are reported.

The research was ethically cleared by Ethical Review Board of Nepal Health Research Council. Prior to participation of student in survey, administrative permission from respective schools and informed consent from student's parent was obtained ensuring voluntary and privacy and confidentiality. Parents and students were also informed beforehand of the study and written informed consent was obtained.

## Results

Around 4.57 % of research participants had went hungry in last 30 days. Similarly, 4.36% had anxiety and 6.27% had felt lonely. Almost two third (65.77%) of the research participants had at least 3 friends. Slightly more than half (50.72%) had experienced bullying in school, 40.09% had experienced physical attack and 39.15% had been involved in physical fighting. Similarly, 6.17% had been using tobacco products and 8.58% had already initiated drug use. Around 13.59% had considered suicide while 10.33% had attempted suicide. Around 23.72% of research participants were of age 16 years or older. (Table 2)

**Table 2:**
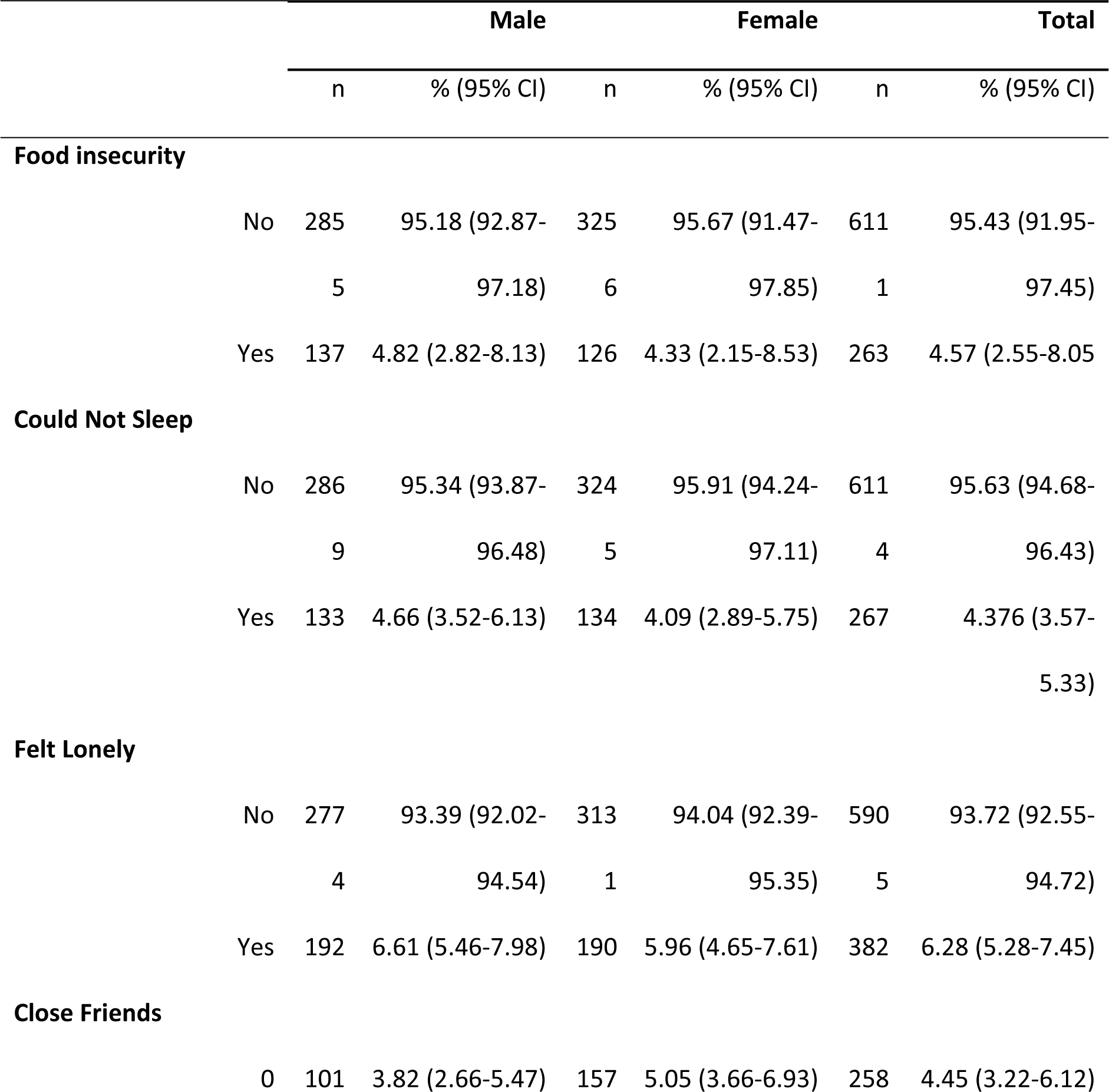
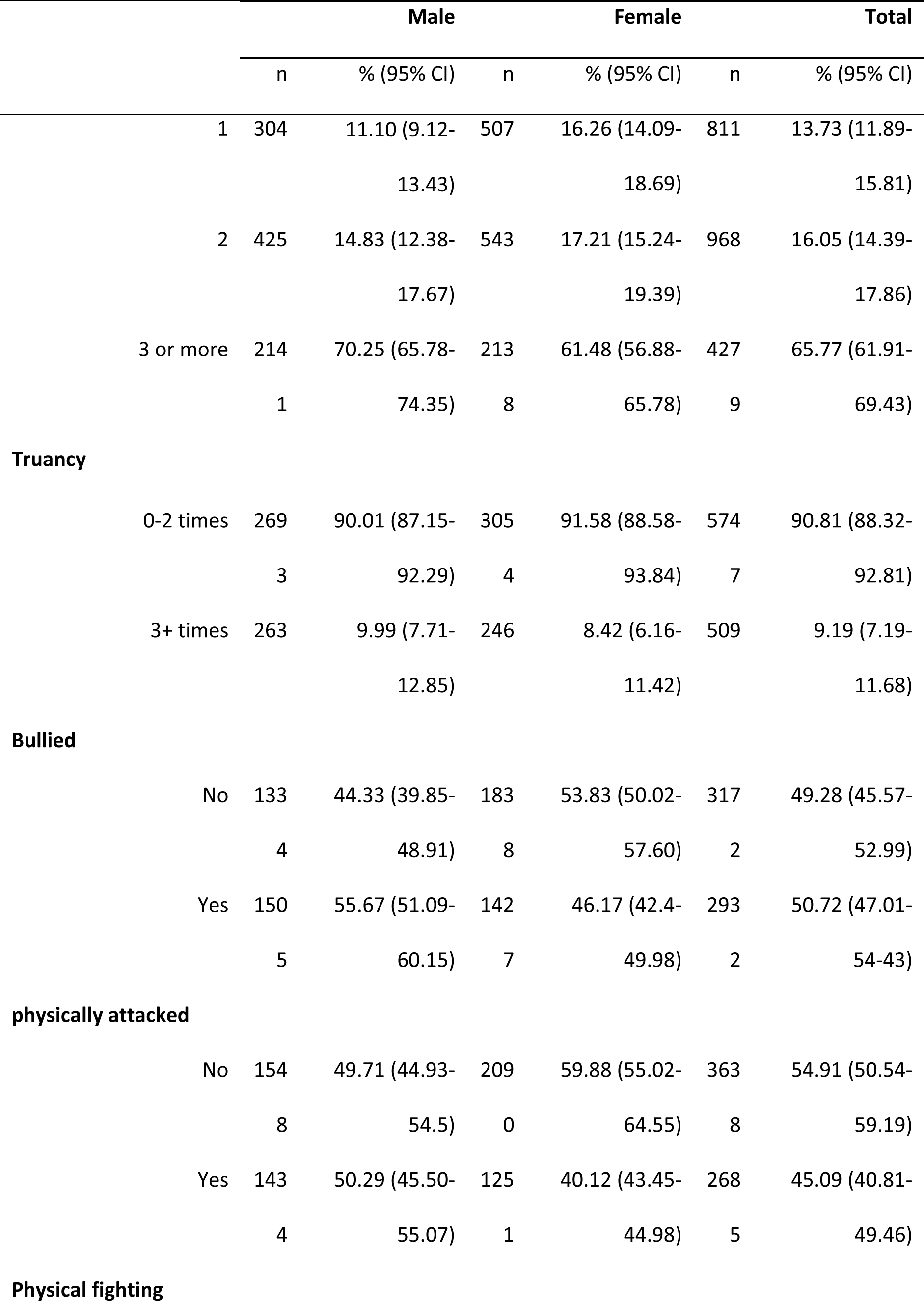
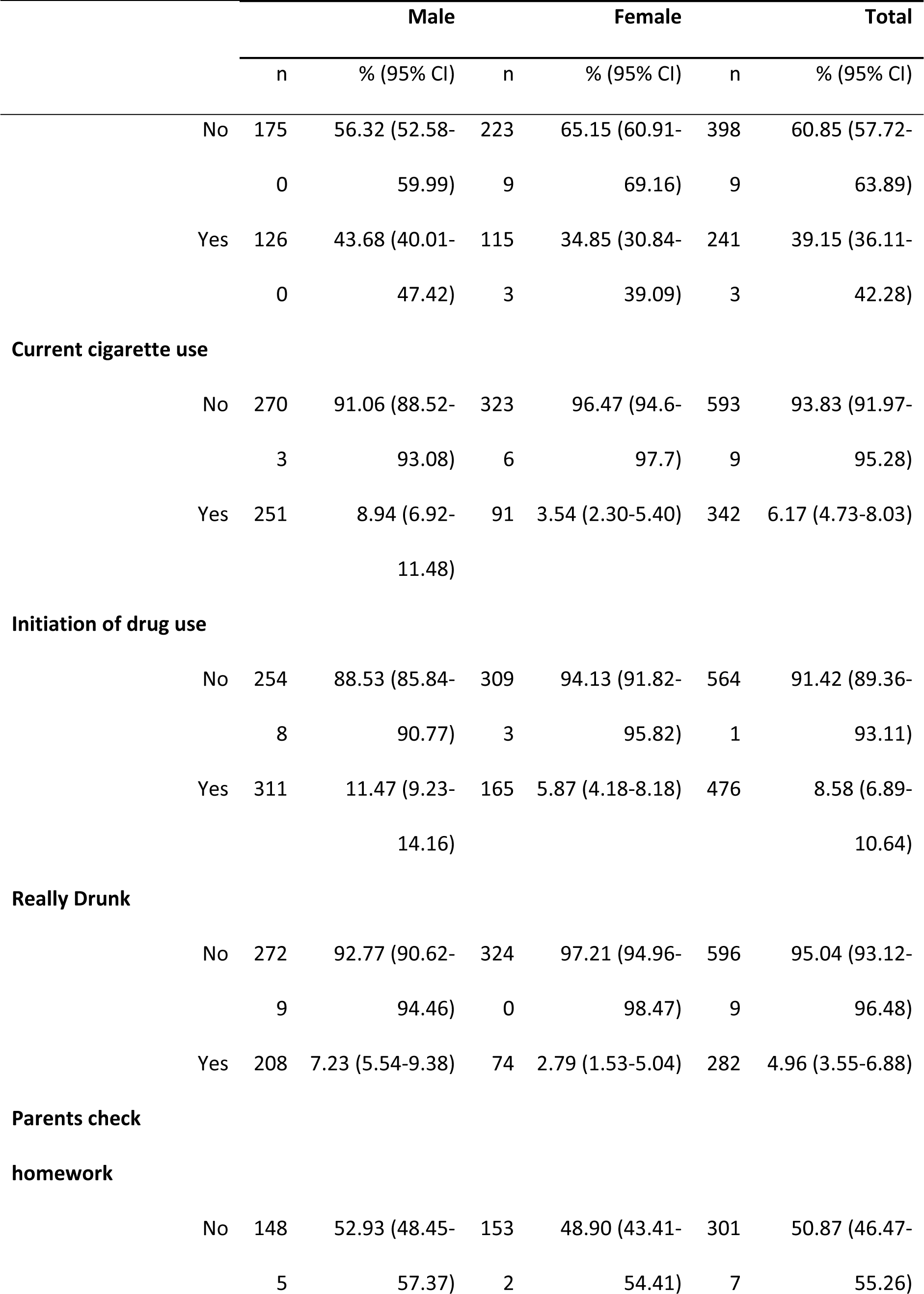
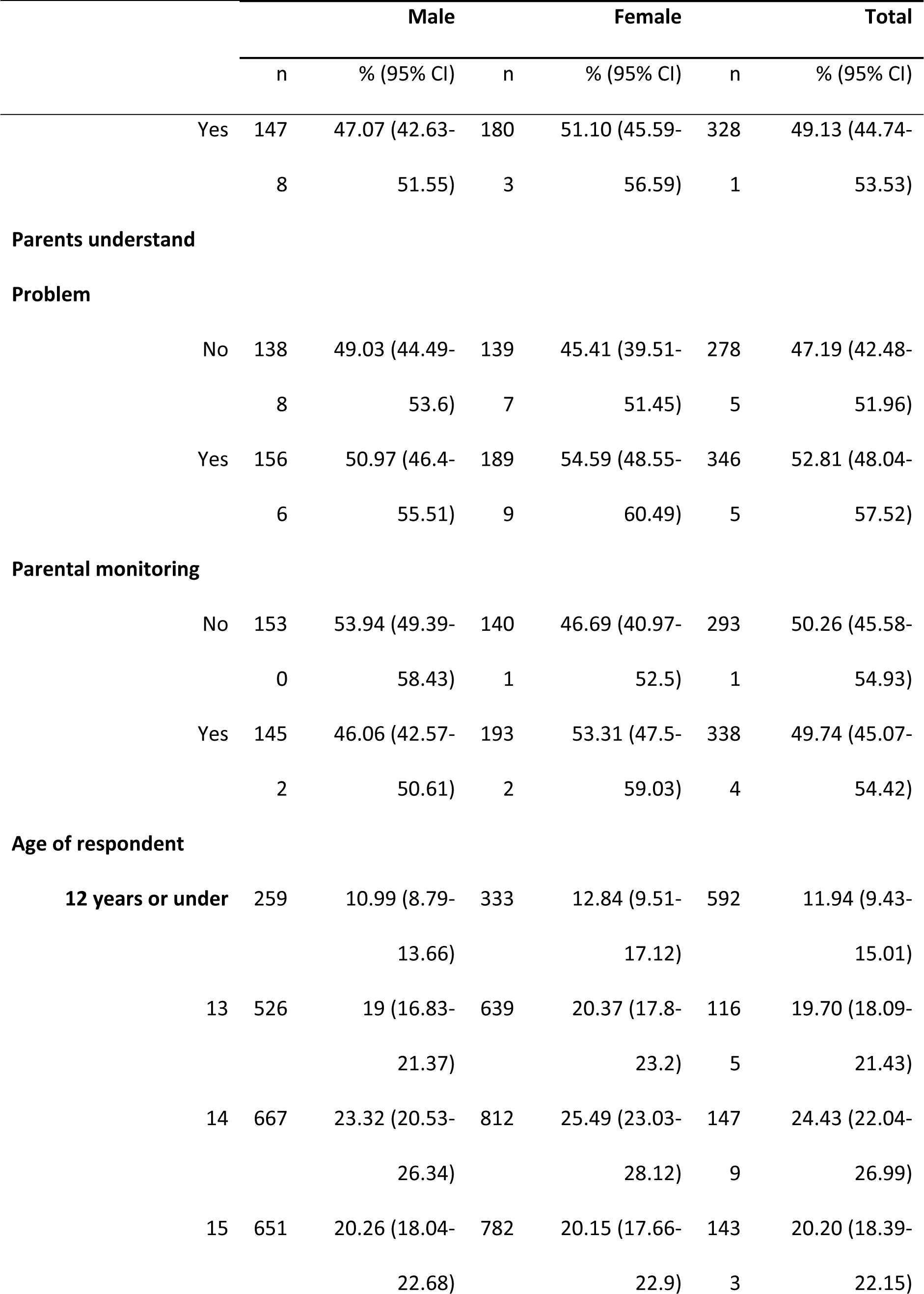
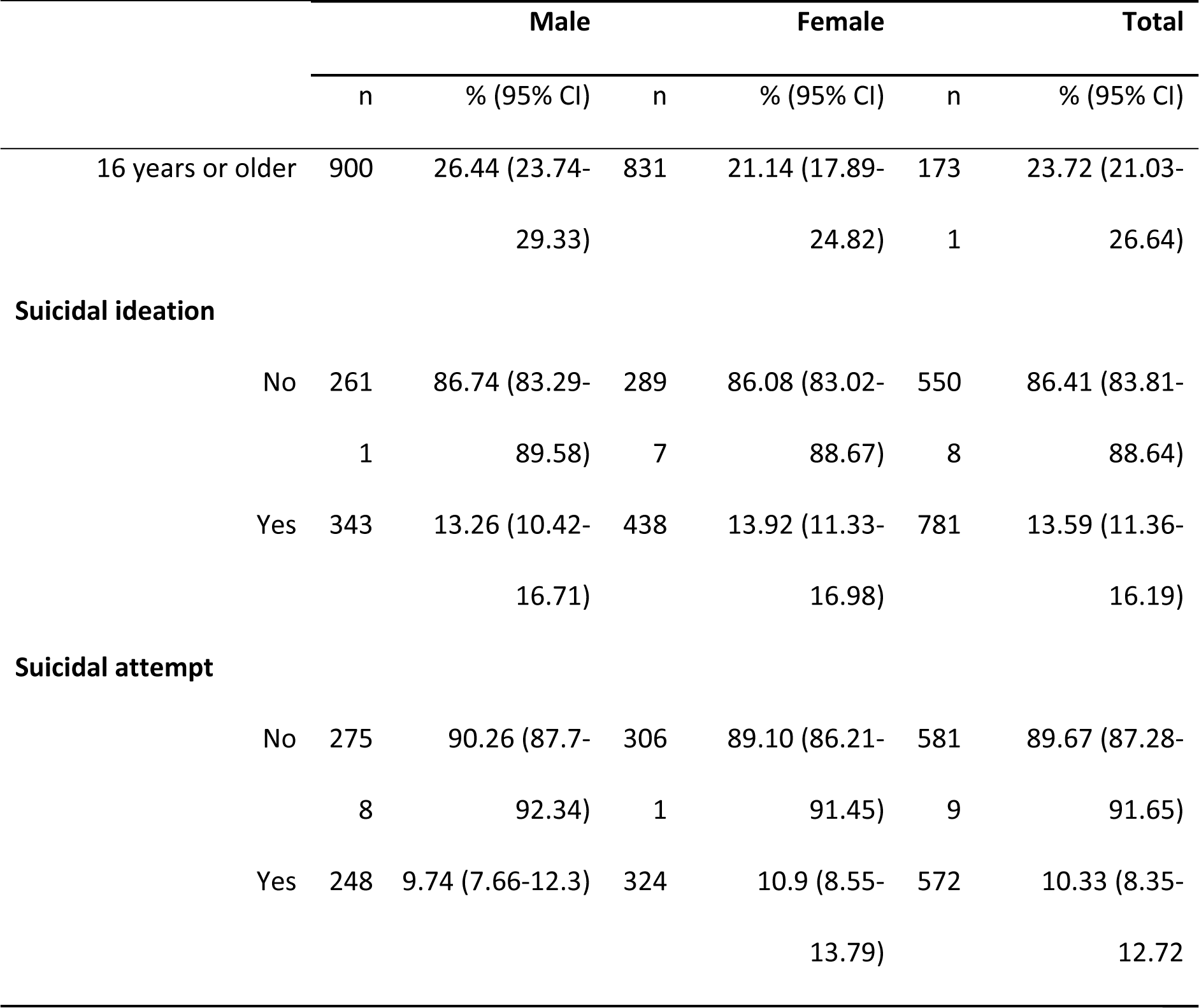
Characteristics of research participants

Adolescents having food insecurity in last 12 months were at 2 folds (OR=2.31, CI=1.64-3.27) higher risk of having suicidal ideation compared to those who had food security. Similarly, adolescents having anxiety had higher odds (OR=2.53, CI=1.48-4.30) of having suicidal ideation compared to their counterparts. Children who felt lonely had higher odds (OR=2.50, CI=1.41-4.44) of having suicidal ideation compared to their counterparts. Similarly, children who had initiated drug use were almost 2 folds (OR=1.61, CI=1.15-2.25) more likely to have suicidal ideation. Similarly, females had higher odds (OR=1.38, CI=1.03, 1.88) of having suicidal ideation compared to males.

Other variables like having parental support, having close friends, missed school, physical fighting, current cigarettes use, getting really drunk, having parents who understand problems and check their homework were not found to have statistically significant association with suicidal ideation. (Table 3)

**Table 3:**
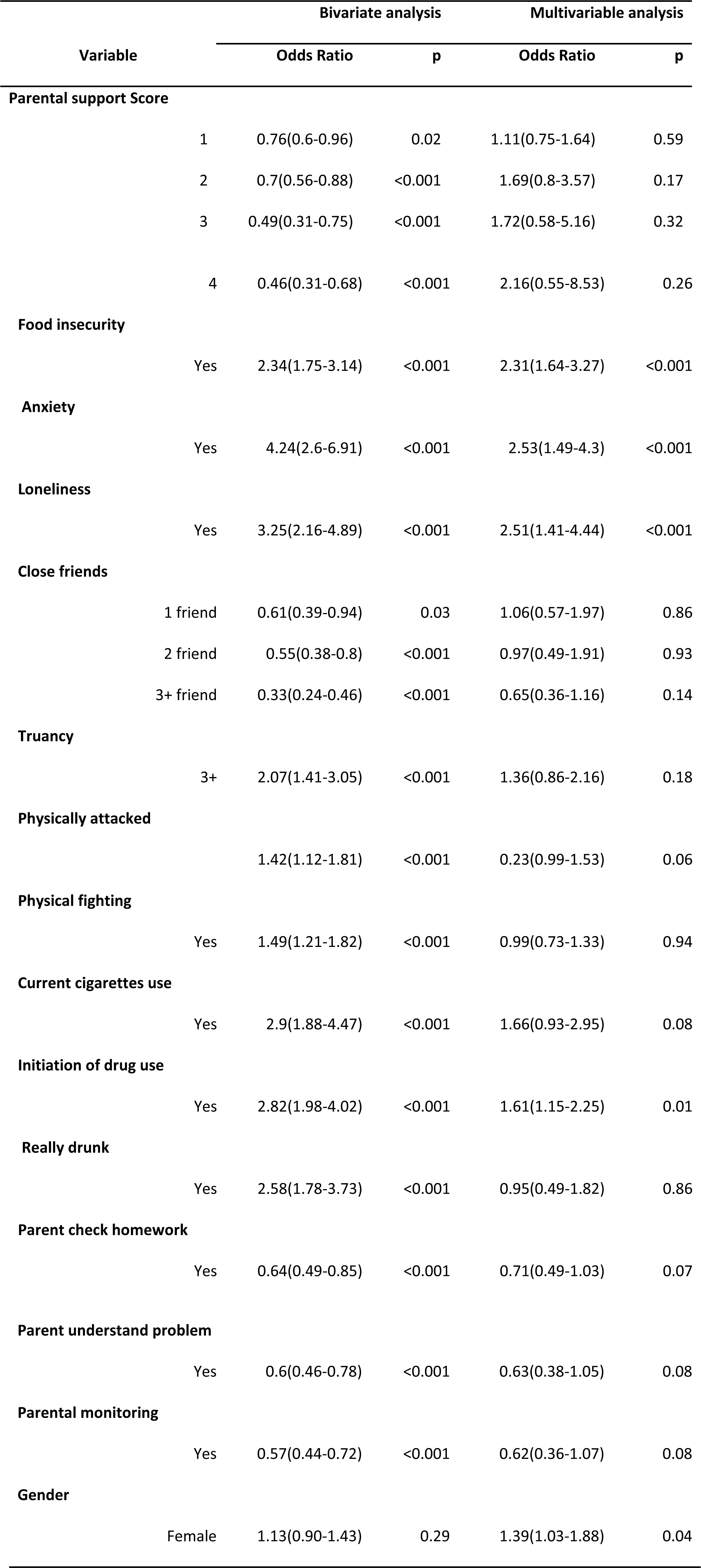
Factors associated with suicidal ideation

| Variable | Bivariate analysis |  | Multivariable analysis |  |
| --- | --- | --- | --- | --- |
|  | Odds Ratio | p | Odds Ratio | p |
| **Parental support Score** |  |  |  |  |
| 1 | 0.76(0.6-0.96) | 0.02 | 1.11(0.75-1.64) | 0.59 |
| 2 | 0.7(0.56-0.88) | <0.001 | 1.69(0.8-3.57) | 0.17 |
| 3 | 0.49(0.31-0.75) | <0.001 | 1.72(0.58-5.16) | 0.32 |

|  |  |  |  |  |  |
| --- | --- | --- | --- | --- | --- |
|  | 4 | 0.46(0.31-0.68) | <0.001 | 2.16(0.55-8.53) | 0.26 |
| **Food insecurity** |  |  |  |  |  |
|  | Yes | 2.34(1.75-3.14) | <0.001 | 2.31(1.64-3.27) | <0.001 |
| **Anxiety** |  |  |  |  |  |
|  | Yes | 4.24(2.6-6.91) | <0.001 | 2.53(1.49-4.3) | <0.001 |
| **Loneliness** |  |  |  |  |  |
|  | Yes | 3.25(2.16-4.89) | <0.001 | 2.51(1.41-4.44) | <0.001 |
| **Close friends** |  |  |  |  |  |
|  | 1 friend | 0.61(0.39-0.94) | 0.03 | 1.06(0.57-1.97) | 0.86 |
|  | 2 friend | 0.55(0.38-0.8) | <0.001 | 0.97(0.49-1.91) | 0.93 |
|  | 3+ friend | 0.33(0.24-0.46) | <0.001 | 0.65(0.36-1.16) | 0.14 |
| **Truancy** |  |  |  |  |  |
|  | 3+ | 2.07(1.41-3.05) | <0.001 | 1.36(0.86-2.16) | 0.18 |
| **Physically attacked** |  |  |  |  |  |
|  |  | 1.42(1.12-1.81) | <0.001 | 0.23(0.99-1.53) | 0.06 |
| **Physical fighting** |  |  |  |  |  |
|  | Yes | 1.49(1.21-1.82) | <0.001 | 0.99(0.73-1.33) | 0.94 |
| **Current cigarettes use** |  |  |  |  |  |
|  | Yes | 2.9(1.88-4.47) | <0.001 | 1.66(0.93-2.95) | 0.08 |
| **Initiation of drug use** |  |  |  |  |  |
|  | Yes | 2.82(1.98-4.02) | <0.001 | 1.61(1.15-2.25) | 0.01 |
| **Really drunk** |  |  |  |  |  |
|  | Yes | 2.58(1.78-3.73) | <0.001 | 0.95(0.49-1.82) | 0.86 |
| **Parent check homework** |  |  |  |  |  |
|  | Yes | 0.64(0.49-0.85) | <0.001 | 0.71(0.49-1.03) | 0.07 |
| **Parent understand problem** |  |  |  |  |
| Yes | 0.6(0.46-0.78) | <0.001 | 0.63(0.38-1.05) | 0.08 |
| **Parental monitoring** |  |  |  |  |
| Yes | 0.57(0.44-0.72) | <0.001 | 0.62(0.36-1.07) | 0.08 |
| **Gender** |  |  |  |  |
| Female | 1.13(0.90-1.43) | 0.29 | 1.39(1.03-1.88) | 0.04 |

Those adolescents anxiety in last 12 months were almost 3 folds (OR=2.99, CI=1.18-7.64) more likely to attempt suicide. Those who felt lonely most of times were almost 2 folds (OR=2.19, CI=1.28-3.74) more likely to attempt suicide compared to their counterparts. Having 3 or more close friends had protective effect (OR=0.35, CI=0.16-0.74) against suicidal attempts although having one or two close friends did not differ significantly from those not having any close friends. Similarly, truancy increased the risk of suicidal attempt by almost 2 folds (OR= 1.99, CI=1.39-2.87). Similarly, those currently using cigarettes were 3 folds more likely (OR=3.06, CI=1.32-7.09) to attempts suicide while girls were around 2 folds more likely (OR=1.60, CI=1.07-2.39) to attempt suicide compared to their counterparts.

Other variables like having parental support, going hungry not having enough food to eat, physical fighting, initiation of drug use, getting really drunk, having parents who check their homework or understand problems were not found to have any statistically significant association with suicidal attempt. (Table 4)

**Table 4:**
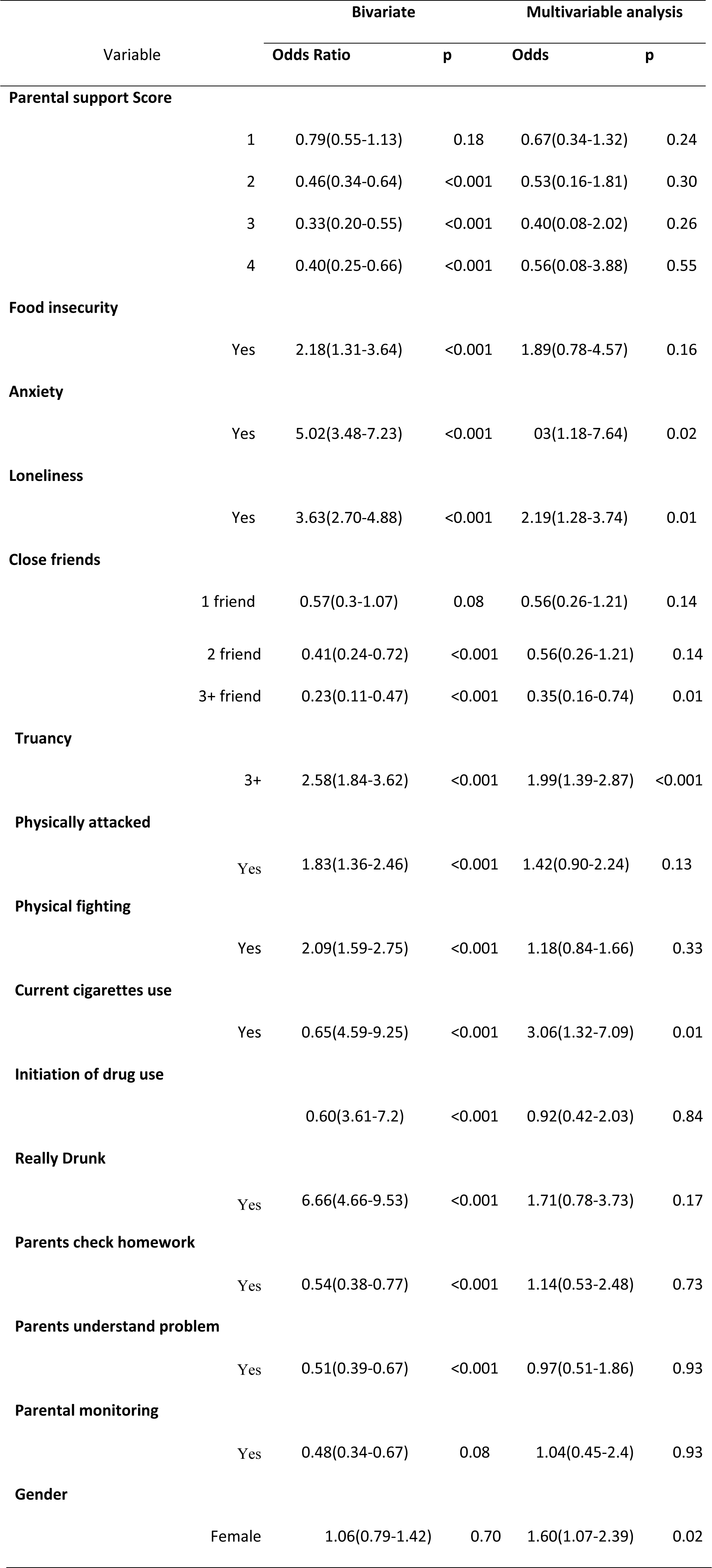
Factors associated with suicidal attempt

| Variable | Bivariate |  | Multivariable analysis |  |
| --- | --- | --- | --- | --- |
|  | Odds Ratio | p | Odds | p |
| **Parental support Score** |  |  |  |  |
| 1 | 0.79(0.55-1.13) | 0.18 | 0.67(0.34-1.32) | 0.24 |
| 2 | 0.46(0.34-0.64) | <0.001 | 0.53(0.16-1.81) | 0.30 |
| 3 | 0.33(0.20-0.55) | <0.001 | 0.40(0.08-2.02) | 0.26 |
| 4 | 0.40(0.25-0.66) | <0.001 | 0.56(0.08-3.88) | 0.55 |
| **Food insecurity** |  |  |  |  |
| Yes | 2.18(1.31-3.64) | <0.001 | 1.89(0.78-4.57) | 0.16 |
| **Anxiety** |  |  |  |  |
| Yes | 5.02(3.48-7.23) | <0.001 | 03(1.18-7.64) | 0.02 |
| **Loneliness** |  |  |  |  |
| Yes | 3.63(2.70-4.88) | <0.001 | 2.19(1.28-3.74) | 0.01 |
| **Close friends** |  |  |  |  |
| 1 friend | 0.57(0.3-1.07) | 0.08 | 0.56(0.26-1.21) | 0.14 |

|  |  |  |  |  |  |
| --- | --- | --- | --- | --- | --- |
|  | 2 friend | 0.41(0.24-0.72) | <0.001 | 0.56(0.26-1.21) | 0.14 |
|  | 3+ friend | 0.23(0.11-0.47) | <0.001 | 0.35(0.16-0.74) | 0.01 |
| **Truancy** |  |  |  |  |  |
|  | 3+ | 2.58(1.84-3.62) | <0.001 | 1.99(1.39-2.87) | <0.001 |
| **Physically attacked** |  |  |  |  |  |
|  | Yes | 1.83(1.36-2.46) | <0.001 | 1.42(0.90-2.24) | 0.13 |
| **Physical fighting** |  |  |  |  |  |
|  | Yes | 2.09(1.59-2.75) | <0.001 | 1.18(0.84-1.66) | 0.33 |
| **Current cigarettes use** |  |  |  |  |  |
|  | Yes | 0.65(4.59-9.25) | <0.001 | 3.06(1.32-7.09) | 0.01 |
| **Initiation of drug use** |  |  |  |  |  |
|  |  | 0.60(3.61-7.2) | <0.001 | 0.92(0.42-2.03) | 0.84 |
| **Really Drunk** |  |  |  |  |  |
|  | Yes | 6.66(4.66-9.53) | <0.001 | 1.71(0.78-3.73) | 0.17 |
| **Parents check homework** |  |  |  |  |  |
|  | Yes | 0.54(0.38-0.77) | <0.001 | 1.14(0.53-2.48) | 0.73 |
| **Parents understand problem** |  |  |  |  |  |
|  | Yes | 0.51(0.39-0.67) | <0.001 | 0.97(0.51-1.86) | 0.93 |
| **Parental monitoring** |  |  |  |  |  |
|  | Yes | 0.48(0.34-0.67) | 0.08 | 1.04(0.45-2.4) | 0.93 |
| **Gender** |  |  |  |  |  |
|  | Female | 1.06(0.79-1.42) | 0.70 | 1.60(1.07-2.39) | 0.02 |

## Discussion

This study identified high burden suicidal ideation and suicidal attempts’ among adolescent students in Nepal. The study has also disentangled the influence of gender, loneliness, having close friend, anxiety, getting drunk, substance abuse, physical fighting and parental support over suicidal ideation and suicidal attempts. This was the first nationally representative study that estimated suicidal ideation and suicidal attempts in Nepalese adolescent students. Moreover, GSHS is conducted in multiple countries among nationally representative samples 13–15-year-olds using globally standardized methodology thus making data comparable from one country to another and provide important opportunity to rectify gap in data on suicide [15]. However, the original study was not designed to determine the factors associated with suicidal ideation and suicidal attempts. Therefore, the factors identified in the study might not fully explain the suicidal ideation and suicidal attempts in the study population because of lacking information on key explanatory variables such as socio-economic status and psychological comorbidities.

Around 13.59% of the research participants had suicidal ideation and 10.3% had attempted suicide in our study. One of the previous study done in four cities in China (Wuhan, Urumqi, Beijing, Hangzhou) had reported that 17.4% of research participants had suicidal ideation and 8.1% had attempted suicide [8]. However, the other study in rural parts of China had found that 19% of the participants had suicide ideation and 7% had attempted suicide attempts in the past year [16]. Prevalence of suicidal ideation was 4.9% in Bangladesh, 11.6% in Bhutan, 13.1% in Maldives, 9.4% in Myanmar, 5.4% in Indonesia, 9.4% in Srilanka, 12.5% in Thailand and 9.3% in Timor-Leste [2,17,18,19,20,21,22,23,24]. GSHS had revealed that suicidal attempt was 6.7% in Bangladesh, 11.3% in Bhutan, 12.7% in Maldives, 8.8% in Myanmar, 3.9% in Indonesia, 6.8% in Srilanka, 13.3% in Thailand and 9.5% in Timor Leste [17,18,19,20,21,22,23,24].

Suicidal ideation and attempt rate seem to differ across countries. Suicidal behavior has a large number of underlying causes. The factors that place individuals at risk for suicide are complex and interactive [8]. Furthermore, access to different means of committing suicide might have created these differences in suicidal ideation and suicidal attempts in different countries.

Our study revealed that girls are at higher risk of suicidal ideation and suicidal attempt compared to boys. Findings from other studies regarding gender differences in suicidal ideation and attempt are not consistent. Most studies previous studies have shown gender differences in suicidal ideation. However, findings differ on whether boys or girls are at higher risk of ideation. Some studies conducted in Canada, Uganda have shown higher rate of suicidal ideation among boys while other studies Malaysia, China and Guyana have shown higher rates among girls [8,25,26,27,28,29]. On the other side, studies conducted in Lebanon, Tanzania, and Thailand have not shown any statistically significant association of suicidal ideation between girls and boys [15,30,31]. Inconsistencies in gender differences in suicidal ideation and attempt might be to social and cultural context of the country that defines status of girls in society. In case of Nepal, being male dominated society, problems of girls might have received less attention that motivated them to consider suicide. Furthermore, in some of the above studies the suicidal ideation might have been underreported by girls thereby showing lower suicide rate [15].

Our study revealed that children who felt lonely most of time or always are more likely to have suicidal ideation and suicidal attempt suicide compared to their counterparts. Findings on association of loneliness with suicidal attempt and ideation are largely consistent in most of previous studies. Previous studies done in Lebanon, Uganda, Tanzania, Sub-Saharan Africa have also revealed that children are more likely to have suicidal ideation when they have feeling of loneliness [15,26,30,32]. Feeling lonely could exacerbate the ill effect of other problems associated with suicide behavior as they find no one to share about the problem that substantially alleviate the agony. This further supported by another findings of our study that having 3 or more close friends had protective effect against suicidal attempts although having one or two close friends did not differ significantly from those not having any close friends. However, its equally important to note that our study revealed no statistically significant association between having close friends and having suicidal ideation. One of the previous study from China had also revealed statistically significant association between having close friends and suicidal attempt and failed to demonstrate any association with suicidal ideation [33]. Another study in Guyana has revealed lower risk of having suicidal ideation if the research participants have close friends [29]. This reinforces the importance of social and peer support in the role of maintaining mental well-being

Similarly, Children who were so worried that they could not sleep most of time in night in past 12 months had higher odds of having suicidal ideation and suicidal attempt. Findings are consistent with most of other studies done in other parts of world. Higher odds of having suicidal ideation was reported in previous studies done in Uganda, Lebanon, Thailand and Republic of Benin when children were worried [15,26,31,34]. Findings are quite usual as adolescents often consider suicide as means to overcome anxiety or distress in life.

There was no significant association between missing school and having suicidal ideation in our study. However, the children who missed school at least 3 times without any permission were found to have almost 2 folds higher risk of having suicidal attempt. Previous similar study done in Republic of Benin had not found any significant association of truancy with suicidal ideation and attempts [34]. However, the variable should be dealt in conjugation with other factors like feeling lonely, having close friends and being worried.

Cigarettes smoking seem to increase the risk of suicidal attempt by almost 3 folds in our study. Some of the previous studies have also demonstrated the association between suicidal behavior and smoking [35,36,37,38,39]. However, there have been some possible explanations for association between smoking and suicide. Suicide has been linked with depression as people might tend to self-medicate themselves with nicotine opting for cigarette smoking or smoking can be personality characteristics associated with low self-esteem [40,41]. Depression might have worked as residual confounder in this study distorting the association of smoking and suicide. One of the previous study also suggests that the association between suicidal behavior and smoking might be because of some unobserved background variables like life circumstances. In the particular study, the use of fixed-effects regression models controlling for unobserved confounding sources had substantially reduced the magnitude associations between suicidal behavior and smoking [42]. Given the lack of any direct plausible causative mechanism, the interpretation of linkage between smoking and suicide need some precautions [43].

There are some limitations in the study. The existing stigma surrounding suicidal ideation and attempts in culturally diversified Nepalese societies might have caused an underreporting of the conditions because of social desirability bias. Study does not cover the adolescent who did not attend school. Furthermore, study also did not collect information on socioeconomic status, religious affiliation, social participation and psychological co-morbidities that could be important in characterizing suicidal ideation and attempt. Further research could be useful in this regard. Despite these limitations, this is the first study done in Nepal among large and nationwide representative sample intended to determine the risk factors of suicidal ideation and attempt. Furthermore, since the study has used globally standardized methodology of global school health survey, study findings are also comparable to other countries adopting same or similar methodology. The study comes out with nationwide estimate of suicidal ideation and attempt among school going adolescents. The study findings could be useful for policy makers in designing appropriate preventive strategies for suicide prevention programs. Adopting appropriate preventive strategies could be very useful in context of Nepal considering the limited availability of treatment services in rural areas constrained with lack of financial and appropriate human resources for mental health services.

## Conclusions

Study reveals high suicide rate among Nepalese school going adolescents. Factors like food insecurity, anxiety, Loneliness and gender were found to be associated with suicidal ideation while anxiety, loneliness, truancy, cigarette use and gender were found to be associated with suicidal attempt.

